# COGEM: A Toolbox for Computational Genomics in Matlab

**DOI:** 10.1101/174896

**Authors:** Theodore J. Perkins

**Author notes:** Web: www.perkinslab.ca.

## Abstract

**Motivation:** The Matlab programming language is widely used for both teaching and research in engineering, computer science, and mathematics. Despite its many strengths, it has never been a dominant language in computational genomics or bioinformatics more generally.

**Results:** Here, we introduce COGEM, a long-term project to develop computational genomics functionality in Matlab. The initial release provides functions for manipulating genomic intervals, stranded or unstranded, with or without numerical data associated. It includes features for both text and binary file input and output, conversion between BAM, BED and BEDGRAPH formats, and numerous functions for manipulating intervals, including shifting, expanding, overlapping, intersecting, unioning, finding nearest intervals, piling up intervals, and performing unary and binary numerical and logical operations on sets of intervals. The toolbox is well-suited to the analysis of high-throughput sequencing data. We demonstrate its functionality by creating a ChIP-seq peak-calling algorithm by chaining together a series of commands, and find it capable of analyzing genome-scale data in reasonable time.

**Availability:** The current toolbox and reference manual is available as supplementary material, and updated versions will be maintained at www.perkinslab.ca online.

## 1 Introduction

Matlab ® (MathWorks, Natick, MA, USA) is a professionally designed and maintained programming language, used widely in teaching engineering, computer science, and mathematics. It has many powerful built in features supporting linear algebra, signal processing, image analysis, machine learning, mathematical programming and visualization, among others. It also has many optimizations allowing for efficient computation, including vectorization, explicitly controllable parallelism, GPU computing, etc. Matlab has become a standard programming language for conducting research in some areas of bioinformatics, including bioimage informatics (Peng, 2008) and systems biology (Schmidt and Jirstrand, 2005; Ghosh *et al.*, 2011). And although some large genomics project support access to their data from Matlab (Cerami *et al.*, 2012), Matlab itself has not become a standard language for conducting computational genomics research.

Our lab has initiated a project to develop, implement and maintain a suite of Matlab functions supporting computational genomics. Our long-term goals are: to facilitate the entry of quantitatively trained students into the area of genomics; to enable cross-fertilization with other areas of bioinformatics (e.g., as genomics and systems biology begin to merge with the rise of single-cell high-throughput sequencing (Wang and Bodovitz, 2010)); and to begin forming a community code base and resource for computational genomics in Matlab, along the lines of the very successful bioconductor effort (Huber *et al.*, 2015). The initial release of our toolbox provides functions for input, output and manipulation of genomic intervals.

## 2 COGEM’s capabilities

The COGEM reference manual (see supplementary information) contains a full description of the software contents. Briefly, there are two main data structures for in-memory representation of genomic intervals: one for BED-formatted data, and one for BEDGRAPH-formatted data (see https://genome.ucsc.edu/FAQ/FAQformat.html for detailed definitions). BED data represents genomic intervals in terms of a chromosome, a start position and an end position. Additionally, strand may be specified, as may be a number of other pieces of information, such as a “name” for the interval, a “score”, etc. COGEM keeps track of all this information, althougth the focus of its manipulations is on the chromosome, start, end and strand information. BEDGRAPH data represents unstranded genomic intervals via a chromosome, start and end position, along with a numeric “dataValue”.

COGEM includes functions for converting from BED to BEDGRAPH (dropping any strand information, and assigning a default dataValue), from BEDGRAPH to BED (dropping dataValue), and from BAM to BED (relying on Matlab’s built-in bamread function; this is useful for converting mapped, high-throughput sequencing reads to BED intervals).

At present, COGEM comprises 34 functions, which enable: reading and writing BED and BEDGRAPH information from and to text files; creating new BED or BEDGRAPH variables; testing whether a set of intervals is sorted, and whether it is disjoint; selecting, shifting, resizing and sorting intervals; finding overlaps between two sets of intervals; finding the nearest intervals in one set to the intervals in another set; coalescing overlapping or adjacent intervals; computing the union and intersection of two interval sets; and performing unary and binary operations on BEDGRAPH intervals (including all the basic arithmetic and logical operators). BED or BEDGRAPH inputs to the functions can be provided either as in-memory variables or as names of files containing the data, and output can be either by in-memory variable or writing to file.

## 3 Demonstration on ChIP-seq peak-calling

As a demonstration, we used the functions in COGEM to develop a simple ChIP-seq peak-calling pipeline intended for single-end reads. It takes as input two files of mapped read locations in BED format, representing ChIP-seq reads and control reads, and outputs a set of peaks with statistically significant enrichment of ChIP-seq reads over control reads. To be clear, our intent is not propose a state-of-theart, competetive peak-calling pipeline, but rather to demonstrate some of the capabilities of COGEM. The 12-step procedure is spelled out in Algorithm 1, with nearly every step being accomplished by a single COGEM command. A Matlab script, which includes time-measurement commands, is available in the supplementary material. Briefly, in the first two steps, mapped read locations are extended in the direction of the strand they are on to an expected read length of 200 (which we assume is independently estimated). Then, the extended ChIP-seq reads are converted to bedgraph format (step 3), where the pileup is computed (step 4) and tested for regions strictly greater than height of 3 (step 5; motivation for threshold of 3 is simply to ignore isolated small read pileups as being certainly without interest). The regions exceeding the threshold are selected (step 6) and coalesced where adjacent (step 7), then converted back to BED intervals (step 8). These are the candidate peaks. ChIP-seq and control reads overlapping those candidate peak regions are counted (steps 9, 10) and statistical significance of the excess of ChIP reads over control reads is evaluated using a Poisson model. Finally, the significant regions are selected and output as peak locations in BED format (step 12). Figure 1A shows the results of several of the steps on a simulated dataset, where the pipeline correctly detects four (obvious) enriched regions. Although the pipeline lacks a few details compared to established peak-callers (e.g., Zhang *et al.* (2008); Rozowsky *et al.* (2009); Zang *et al.* (2009)), we consider it a plausible pipeline that, more importantly, demonstrates the versatility of the functions in the COGEM distribution, and the ease with which they allow rapid prototyping.

**Figure 1:**
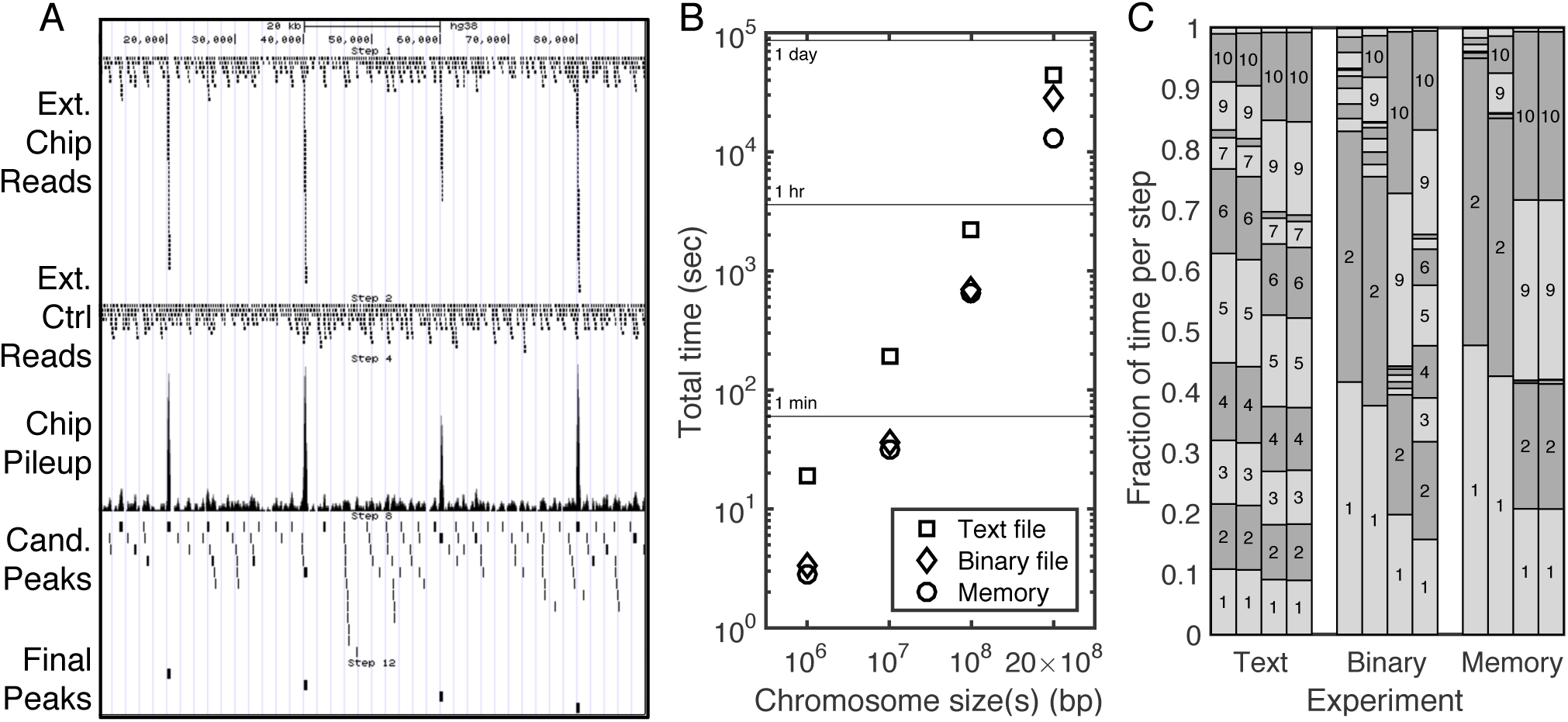
Results of a ChIP-seq peak-calling pipeline developed using the COGEM toolbox. (A) Major steps in the analysis, visualized with the help of the UCSC Genome Browser (Kent *et al.*, 2002). (B) Total time taking by the pipeline, depending on the chromosome/genome size and whether each step is written to and from text file, to and from a binary file, or simply kept in memory. (C) Fraction of time taken by each step. Each group of four columns reports times for the four chromosome/genome sizes tested.

To test the pipeline, we generated simulated ChIP-seq data of various sizes: either 10^4^ reads spread over a 10^6^ bp hypothetical chromosome, 10^5^ reads on a 10^7^ bp chromosome, or 10^6^ reads on a 10^8^ bp chromosome. The smallest chromosome is similar to the size of the whole genome of some smaller bacteria, while the largest chromosome size (100 megabases) is close to the median human chromosome size. At each size, we generated 10 simulated ChIP-seq datasets, in which 80% of the reads were uniformly spread over the chromosome, and 20% of the reads were concentrated in 500bp windows spread approximately 20kb apart. We generated 10 matching control datasets where 100% of reads were uniformly spread. We also generated a simulated ChIP-seq and control dataset comprising 2 *×* 10^7^ reads over a hypothetical genome of 20 chromosomes, each with 10^8^ bp. These choices approximate the size of the human genome and the size of a modern ChIP-seq study.

### Algorithm 1

A simple ChIP-seq peak-calling pipeline constructed using functions from the COGEM toolbox.

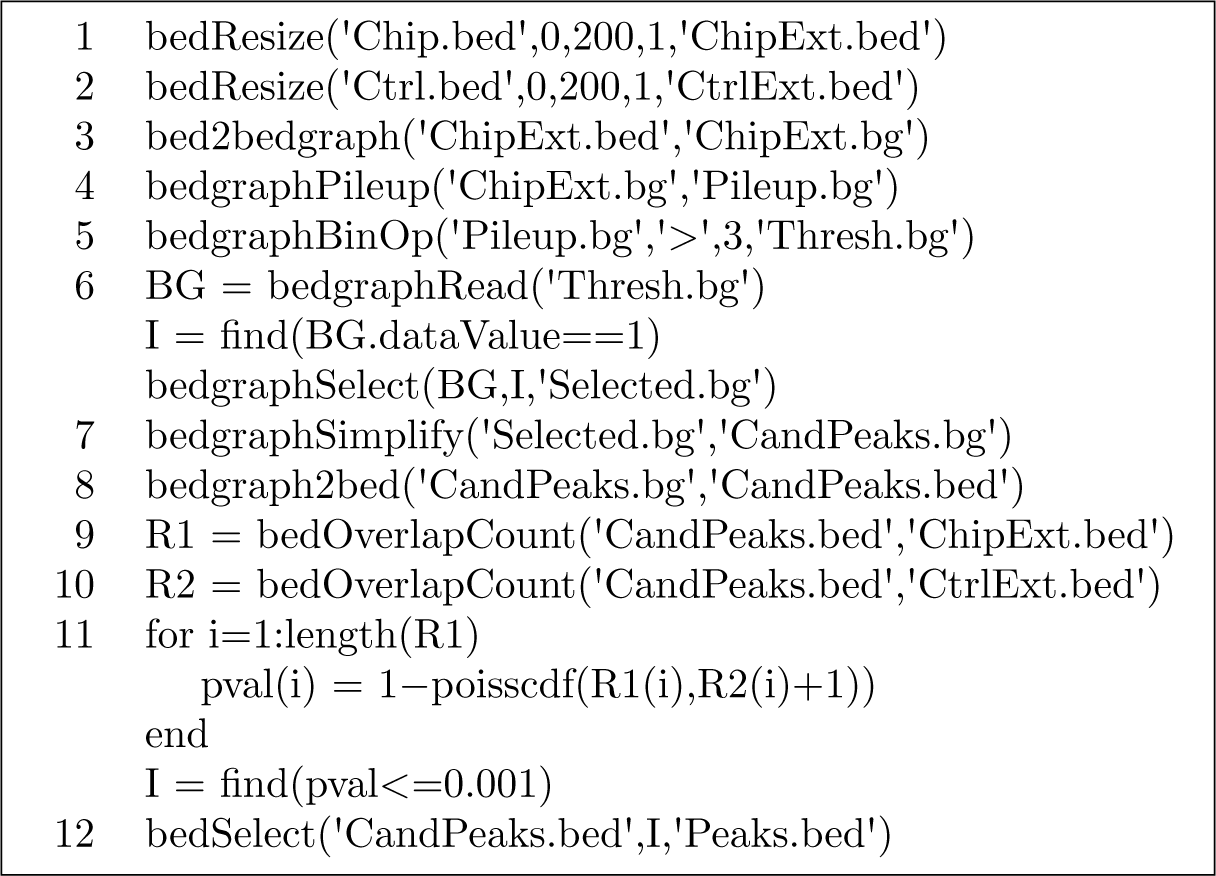

We then ran the pipeline, measuring the execution time of each step on a MacBook Pro with 2.6 GHz Intel Core i7 CPU and 16 GB RAM. The squares in Figure 1B shows the total average run time for the three chromosome sizes and the approximately-human-genome size. The left side of Figure 1C shows the fraction of time spent in each of the twelve steps of the pipeline. The fastest steps were 8 (format conversion for the relatively small number of candidate peaks), 11 (statistical significance, once reads in the candidate peaks have been counted), and 12 (outputting final peaks). The remaining steps all took roughly the same amount of time, and that is largely due to the text-formatted file input and output. We also tested two other variants of the pipeline. In one, results are saved to and re-read from binary files at each step, using the Matlab “save” and “load” commands applied to the in memory BED or BEDGRAPH variable resulting from each step. In the other variant, the only file i/o is the initial BED file input and final peak output; all other steps take place in memory. These results are shown in Figures 1B (diamonds and circles) and 1C (middle and right blocks). For the single chromosome computations, either variant saves nearly an order of magnitude of time. The variability in run times across the 10 random repeats was less than 7% in all cases, and less than 1% in most cases. For the genome-scale computations, the binary-file variant remains faster than the text-file variant, but the in-memory variant is substantially better. The total times for text, binary and memory versions are 12.41 hours, 8.02 hours, and 3.63 hours. For the two smallest dataset types, the binary and in-memory versions of the pipeline spend most of their time on the initial reading in of the BED files. For the larger datasets, initial input remains time consuming, but counting the overlaps of ChIP-seq and control reads with candidate peaks, which is a quadratic operation as currently implemented, also becomes important.

## 4 Conclusion

Computational genomics is a burgeoning field, driven by new technologies creating massive amounts of data. Appropriate software tools are needed to analyze that data. While various tools are available in various languages, pipelines often must be cobbled together using different resources, requiring different formats, etc. Students are usually not trained in all the different languages required. Here, we put forth COGEM, a toolbox supporting computational genomics, written in Matlab—a language that is part of the training of many undergraduate students. It focuses on the manipulation and analysis of genomic intervals, as might arise from high-throughput sequencing data. We showed that with minimal effort, a plausible ChIP-seq peak-calling pipeline could be constructed that runs efficiently enough to be used on modern, genome-scale datasets. Intended future efforts to expand the COGEM toolbox include support for variant calling and detection of other mutational events (useful in personalized medicine and cancer), mutational and expression biomarker identification, co-expression analysis and single-cell sequencing data analysis. Researchers interested in joining the COGEM project to bring computational genomics to Matlab are encouraged to contact the author. Collaboration is welcome.

## Acknowledgements

We thank Julie Fiala and R. Matt Tanner for helping to test the COGEM code and for providing feedback on this manuscript.

### Funding

This work has been supported by the Natural Sciences and Engineering Research Council of Canada (NSERC), grant number 328154.

